# Advection and habitat loss interactively reduce persistence: maintaining threatened riverine populations while restoring natural flow regimes

**DOI:** 10.1101/717496

**Authors:** Vadim A. Karatayev, Lyubov E. Burlakova, Alexander Y. Karatayev, Luojun Yang, Thomas Miller

**Affiliations:** Department of Environmental Science and Policy, University of California, Davis. One Shields Ave Davis, CA 95615.; Great Lakes Center, SUNY Buffalo State; School of Life Sciences, Nanjing University. Present address: Department of Ecology and Evolutionary Biology, Princeton University; Environmental Science Center, Laredo Community College

**Keywords:** Spatial population dynamics, dispersal, catastrophic drift, habitat loss, unionidae, environmental flows

## Abstract

Modification of flow regimes and habitat degradation are the strongest, most common, and often co-occurring human activities affecting riverine populations. Ongoing efforts to restore peak flow events found under pristine flow regimes could increase advection-driven dispersal for many species. In rivers with extensive habitat loss, increased advection could transport individuals from remnant populations into degraded downstream areas, causing restored flow regimes to decrease persistence of threatened species. To resolve the capacity for such ‘washout’ effects across imperiled taxa, we evaluate population growth in spatial matrix models of insect, fish, and mollusc taxa experiencing advective dispersal and either long-term habitat loss or temporary disturbances. As a case study to quantify advective dispersal in threatened species, we use intensive mark-recapture methods in a Rio Grande population of the federally endangered unionid mussel Texas horhshell (*Popenaias popeii*). Among unionids, the most threatened freshwater taxa of North America, we find high levels of annual downstream emigration (16-51%) of adult *P. popeii*, concomitant with strong immigration from upstream habitats. For different taxa experiencing such advective dispersal during specific life stages, our population model shows that washout effects strongly reduce population recovery under high levels of habitat loss. Averting this negative consequence of restoring hydrology requires simultaneously restoring or protecting long, contiguous stretches of suitable habitats. Across taxa in heavily impacted systems, we suggest integrating hydrodynamic studies and field surveys to detect the presence of advective dispersal and prioritize areas for habitat restoration to enhance population persistence.

## Introduction

Steeply rising human impacts over the past century have made freshwaters among the most endangered ecosystems, with a nearly tenfold higher concentration of imperiled species compared to marine and terrestrial habitats (Strayer and Dudgeon 2010). Decline and ultimate extirpation of native species within ecosystems characteristically occurs in response to multiple human-induced stressors (Dudgeon et al. 2006). Emerging conservation approaches optimize dam discharges to mimic natural flow conditions (“environmental flows”, Tharme 2003; Yarnell et al. 2015) while others focus on restoring pristine habitats at smaller scales by altering riparian vegetation, riverbed substrates, or nutrient loading (Bernhardt et al. 2005). However, co-occurrence of multiple human impacts in degraded ecosystems might limit success of efforts focused on alleviating individual stressors (Ormerold et al. 2010).

Flow modification is a ubiquitous anthropogenic impact on rivers (e.g., 85% of US rivers dammed, Poff & Hart 2002; Tharme 2003) that has transformed the water quality, food web energy sources, and physical habitats in lotic ecosystems. Environmental flows aim to help restore such ecosystem functions in part by mimicking the magnitude of extreme discharge levels (Yarnell et al. 2015). Simultaneously, peak flow events predominantly regulate the spatial distribution and mass transport of organisms with river currents (‘advective dispersal’ hereafter), as commonly found in aquatic insects (Brittain & Eikeland 1988; Gibbins et al. 2007), juvenile fish (Lechner et al. 2016), and molluscs (Gangloff & Feminella 2007; Kappes & Haase, 2012). Consequently, implementing environmental flows can intensify advective dispersal in one or more life stages for a broad range of taxa.

In heavily degraded rivers, restoring peak flows present under pristine conditions might interact with habitat loss to accelerate loss of native taxa by consistently moving individuals out of remnant pristine habitats into degraded ‘sink’ areas (‘washout’ effect hereafter; Speirs & Gurney 2011; Lutscher et al. 2006). Loss of natural river habitats typically arises from reservoir construction, water depletion in riverbeds, or when intensified runoff from heavily developed terrestrial landscapes causes eutrophication and siltation (Allan & Flecker 1993). Such stressors typically arise in arid or agricultural regions where dams regulate flow to provide flood control, irrigation, and water storage (Poff & Hart 2002), and only some forms of habitat loss might quickly subside under natural flow regimes. As a result, preserving threatened populations might require restoring riverine habitats and flow regimes simultaneously to avert washout effects.

The capacity of washout effects to undermine population persistence may depend on the magnitude of advective dispersal and the forms of habitat loss. Across taxa, this potential might depend on the life stages vulnerable to advection and their contribution to population growth. Across systems, washout effects may also depend on the spatial scale of habitat loss: for instance, reservoir construction produces large, isolated subpopulations whereas localized streambed channelization or siltation intersperses sink habitats, and subsequent washout effects, throughout the entire population. Within specific taxa, the presence and magnitude of advective dispersal is also rarely resolved, especially among endangered species. For example, freshwater mussels (Unionoida order, >300 mostly river-dwelling species) are considered the most imperiled faunal group in North America (Master et al. 2000), with strong fragmentation and declines of populations primarily linked to long-term habitat loss (Strayer 2008; Downing et al. 2010). While some studies have discussed the presence of advective dispersal in adult unionids (Balfour & Smock 1980; Strayer 1999; Hastie et al. 2001) or demonstrated its effects on the distribution of juveniles (Daraio et al. 2012), most intensive population studies do not consider advective dispersal (e.g., Villella et al. 2004; Matter et al. 2013), even in long-lived populations annually experiencing intense flow events. At same time, washout effects could have strong negative consequences for populations by affecting juvenile or long-lived, critically important adult stages.

Here we analyze the capacity for washout effects to reduce persistence in a wide range of taxa. For this we develop a spatially explicit population model spanning distinct life histories that incorporates advective dispersal and qualitatively different forms of habitat loss. In conjunction, we quantify the extent of advective dispersal in a mussel population subject to extensive habitat loss, the federally endangered Texas hornshell (*Popenaias popeii*) in the Rio Grande, using a large-scale survey and intensive mark-recapture study of population dynamics and demography. We then discuss systems and taxa where washout effects might most strongly affect population persistence, with clear implications for conservation, and evaluate specific approaches to detecting and quantifying advective dispersal across systems.

## Methods

### Spatially structured population model

We developed a general, spatially explicit population model to examine washout effects on population persistence across taxa and conservation scenarios (Fig. 1). We modeled space discretely to capture the spatial clustering seen in most riverine faunal groups among discrete habitat patches, between which animals disperse by advection or actively via swimming, entering river currents, or attaching to other organisms (Townsend 1989). Species within individual patches in turn may undergo either long-term extirpation from habitat loss or temporary extirpation from disturbances like droughts. The scales of habitat patches and habitat loss we considered exceed the distance that individuals vulnerable to advection can actively move. This means that fish larvae, insect larvae, and molluscs transported into degraded patches cannot escape high mortality via swimming or elective downstream drift to better habitats. To resolve how these dynamics impact population recovery or extinction, we analyzed low-density population growth λ in spatial matrix population models (Rogers 1966). These models also readily incorporate the effects of population age or stage structure (Appendix S1).

**Figure 1.**
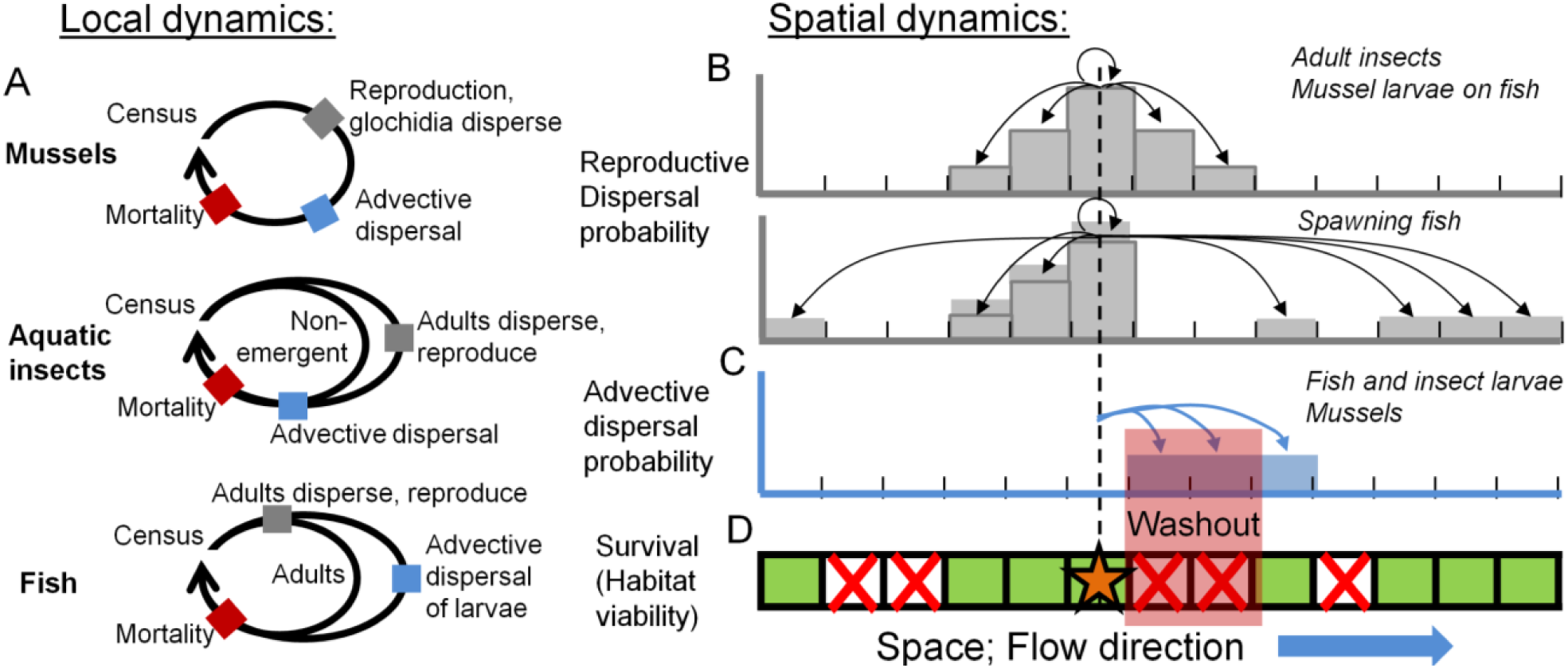
Local dynamics of modeled life histories differ in stages affected by advection (all individuals for mussels and insects versus larval recruits only for fish; a). Across space, recruits from a focal patch are dispersed to adjacent patches (b, upper plot), but fish readily detect and avoid spawning in unviable habitat patches (b, lower plot). Advective dispersal then transports vulnerable organisms downstream (c), and dispersal to unviable habitat patches (red crosses in d) produces washout effects.

Our model tracks N_i,t_, the abundance of adult (sexually mature) individuals in patch i (with patch i = 1 being furthest upstream) in year t, in a population distributed across L habitat patches in a river. Abundance in patch i next year (N_i,t+1_) is the sum of (1) individuals in patch i now (N_i,t_) that remain after natural mortality d and advective dispersal ρ, (2) the number of new recruits produced in surrounding patches (r recruits per adult) and actively dispersed into patch i, and (3) the fraction individuals within m upstream patches which arrive via advective dispersal after surviving dispersal-induced mortality μ. Active dispersal during reproduction spans a mean distance of n patches and is represented by the matrix **F**, wherein 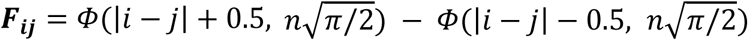 and Φ is the cumulative Gauss distribution (Fig. 1b). Downstream advective dispersal is represented by **D**, wherein **D**_ij_ = (1 -ρ) for i = j, ρ(1 -µ)m^-1^ for 1 ≤ i -j ≤ m, and 0 otherwise (Fig. 1c). Throughout we assume that reproduction occurs before natural mortality and advective dispersal. Finally, the vector 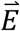 denotes habitat quality of each patch (here 1 for suitable or 0 for unviable habitats), and can vary across years under temporary habitat loss (Fig. 1d).

To examine the effects of habitat loss and advective dispersal on different taxa, we modeled several qualitatively distinct life histories (Fig. 1a). In the case of iteroparous species in which advective dispersal affects both adults and recruits (e.g., freshwater mussels), we modeled adult abundance as

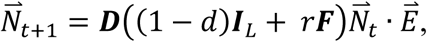

where **I**_L_ is an LxL identity matrix and · denotes the vector dot product. In the case of semelparous aquatic insects our model tracks larvae, of which a fraction γ annually leave the aquatic larval stage to disperse and reproduce as winged adults:

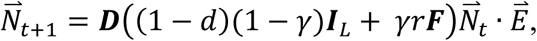

where r is the product of the chance an adult survives to reproduce and the number of larvae they produce. Finally, we modeled fish populations where only recruits experience advective dispersal as larvae, while surviving adults annually move among (and reproduce in) nearby, viable patches. We assumed adults in isolated habitats with no other viable patches nearby disperse to any viable patch in the river by normalizing the columns of 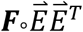 to unity using **M**:

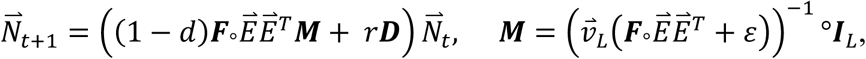

Where ∘ denotes element-wise multiplication, 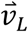 is a length-L unity vector, and ε=10^-9^ is an arbitrary, small constant.

### Washout effects on persistence across taxa

First, we examined the extent to which washout reduces low-density population growth λ across different life histories (Fig. 2). Specifically, we modeled populations of aquatic insects (*Hydropsychidae* sp.) and shorter-*vs.* longer-lived species of fish (*Alburnus alburnus* and *Abramis brama*) and freshwater mussels (*P. popeii* and *Margaritifera margaritifera*). This range of taxa captures life histories differing in the life stage affected by advective dispersal and the relative abundance of recruits at low densities (greater for shorter-lived species). For each life history, we compared the percent decline in population growth λ in the presence of: high habitat loss (85% of patches are unviable), moderate levels of advective dispersal in the vulnerable life stage, and habitat loss and advective dispersal that together produce washout. For all cases, we set per capita reproduction such that, under ideal conditions without advective dispersal or habitat loss (ρ=0, E=0), populations recovering from low densities annually grow by 25% for shorter-lived taxa and by 10% in long-lived species with delayed maturation.

**Figure 2.**
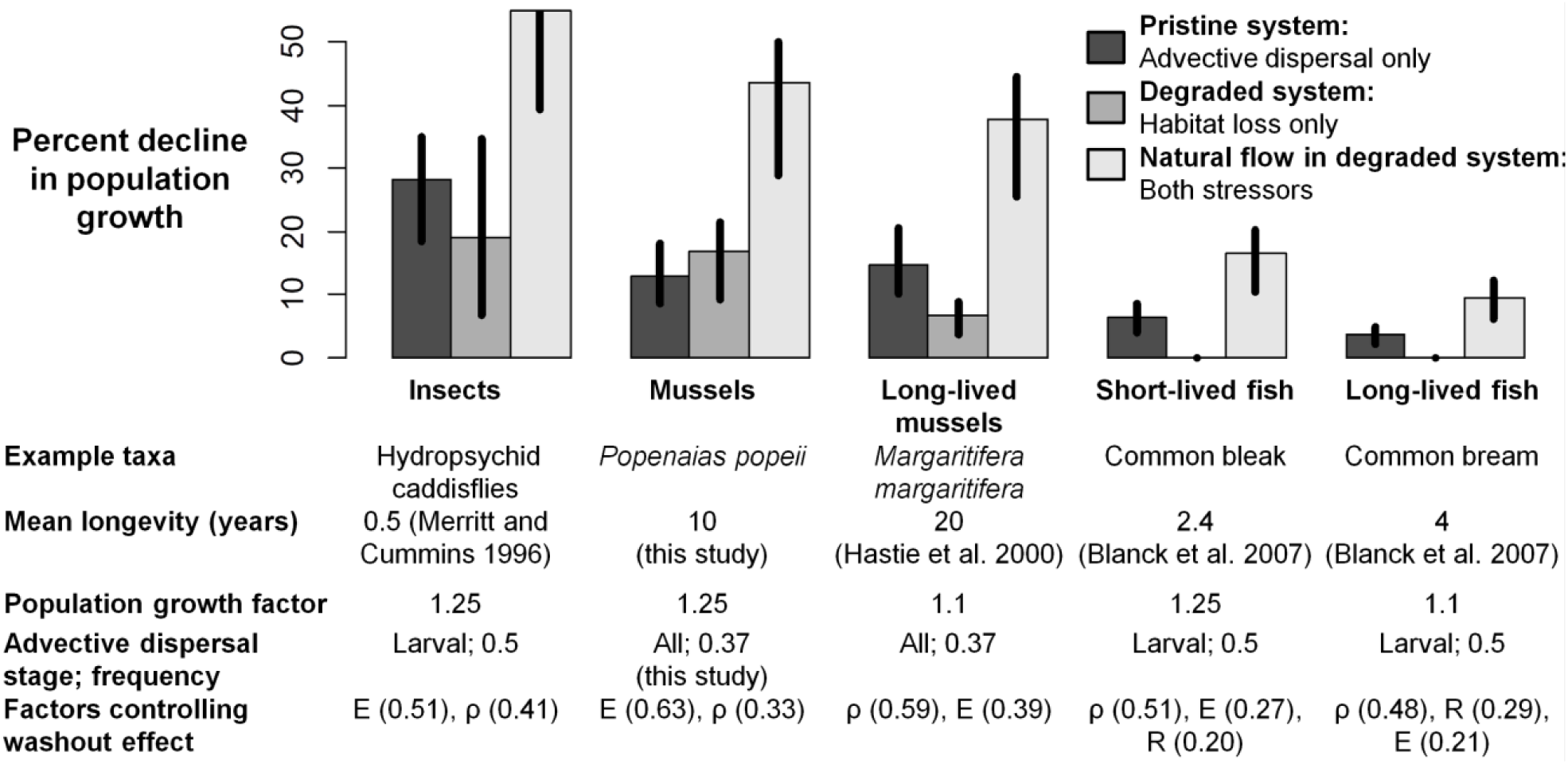
Independent and interactive effects of advective dispersal and permanent habitat loss (85% of patches uninhabitable) on the decline in population growth relative to ideal conditions (i.e., no advective dispersal or habitat loss) across life histories. For caddisfles, mean longevity denotes the average duration of juvenile stages preceding mortality or emergence. Vertical segments denote 25^th^ and 75^th^ percentiles of model results in the global sensitivity analysis varying all model parameters within ±50% of their default values (restricted by y-axis range for insects). Factors controlling washout effect denote parameters having the greatest effect on the decline in growth under both advective dispersal and habitat loss (advective dispersal frequency ρ, extent of habitat loss E, or per capita recruitment R; fractions denote relative importances). See Table 1 and Appendix B for model parameterization.

Throughout, we determined model parameters based on our empirical results (see below), or published studies (Appendix S2). To ensure our results are robust to uncertainty and system-specific variation in parameters, we conducted a global sensitivity analysis (Fig. 2). For this we varied each parameter over a uniform distribution spanning ±50% of the default value, randomly drew 4,000 parameter sets from this multivariate distribution, and for each parameter set re-calculated percent decline in λ under habitat loss, advective dispersal, and both stressors. We then summarized the distribution of these outcomes across parameter sets (vertical bars in Fig. 2). Additionally, we identified parameters that predominantly control washout effects (the decline in λ under habitat loss and advective dispersal) by summarizing our sensitivity results using a random forest approach. This standard sensitivity analysis calculates the importance of each parameter to the model outcome (see Harper et al. 2011).

**Table 1.**
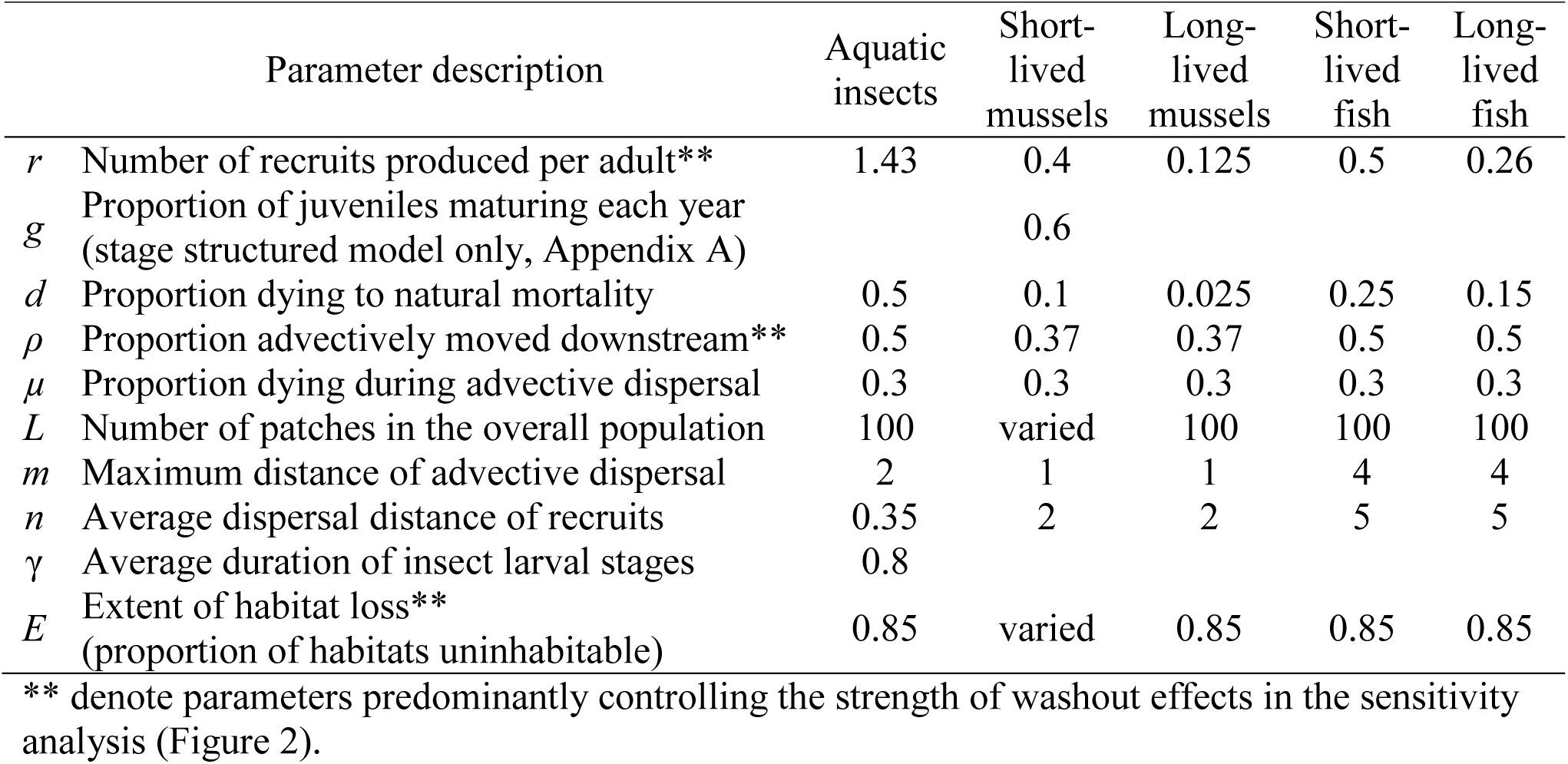
Description and default values of model parameters. Note that dispersal distances are measured relative to the mean distance between adjacent habitat patches. For sources and derivation of parameters see Appendix B.

|  | Parameter description | Aquatic insects | Short-lived mussels | Long-lived mussels | Short-lived fish | Long-lived fish |
| --- | --- | --- | --- | --- | --- | --- |
| $r$ | Number of recruits produced per adult** | 1.43 | 0.4 | 0.125 | 0.5 | 0.26 |
| $g$ | Proportion of juveniles maturing each year (stage structured model only, Appendix A) | | 0.6 | | | |
| $d$ | Proportion dying to natural mortality | 0.5 | 0.1 | 0.025 | 0.25 | 0.15 |
| $\rho$ | Proportion advectively moved downstream** | 0.5 | 0.37 | 0.37 | 0.5 | 0.5 |
| $\mu$ | Proportion dying during advective dispersal | 0.3 | 0.3 | 0.3 | 0.3 | 0.3 |
| $L$ | Number of patches in the overall population | 100 | varied | 100 | 100 | 100 |
| $m$ | Maximum distance of advective dispersal | 2 | 1 | 1 | 4 | 4 |
| $n$ | Average dispersal distance of recruits | 0.35 | 2 | 2 | 5 | 5 |
| $\gamma$ | Average duration of insect larval stages | 0.8 | | | | |
| $E$ | Extent of habitat loss** (proportion of habitats uninhabitable) | 0.85 | varied | 0.85 | 0.85 | 0.85 |

\*\* denote parameters predominantly controlling the strength of washout effects in the sensitivity analysis (Figure 2).

### Washout effects across habitat loss scenarios

We explored how washout effects on persistence depend on qualitatively different forms of habitat loss. In the case where habitat loss happens locally, we examined a system of L=100 habitat patches and vary the proportion of randomly selected patches rendered unviable (i.e., where mortality d=100%; Fig. 3a). Next, we considered large-scale habitat loss (e.g., reservoir construction or intense pollution) that produces demographically isolated populations. For this we examined how the number of adjacent viable remnant habitat patches L affects λ across levels of advective dispersal (Fig. 3b). Finally, we examined washout effects on riverine metapopulations wherein the location of habitat loss (i.e., local disturbances) varies among years (Fig. 3c). We assumed a fraction (30%) of patches is protected from disturbance, as in the case where deeper pools covering a fraction of the riverbed maintain water during drought (Bennett and Bouwes 2009); we also considered the simpler case with all patches equally vulnerable (Appendix C). For both disturbance patterns we used stochastic implementations of our model and calculated λ as the mean of annual population growth factors over 100 years. Throughout, we parameterized the model based on our case study of *P. popeii* in the Rio Grande, but note that qualitatively similar mechanisms and results also occur across the different life histories examined above.

**Figure 3.**
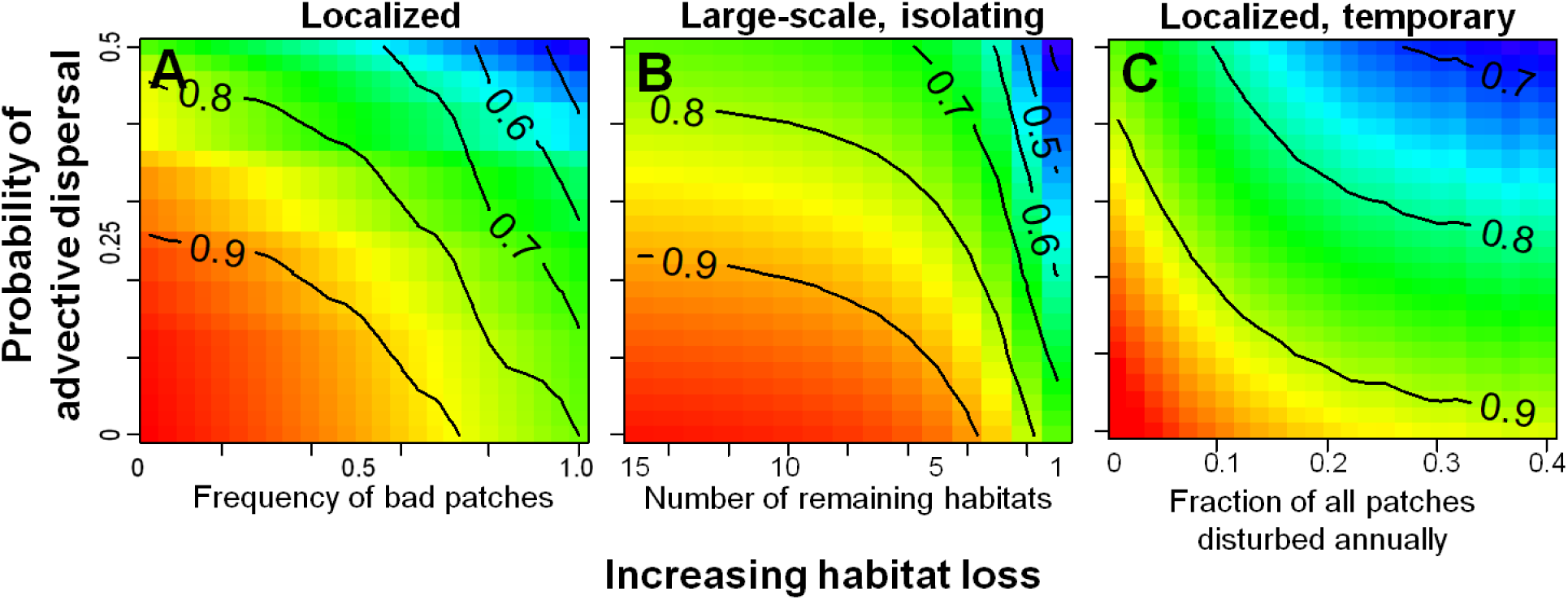
Due to washout effects, advective dispersal (ρ, y-axes) reduces population growth more strongly at high levels of habitat loss (x-axes) that happens either locally in individual patches throughout the system (a), at large scales by reducing the number of remaining patches in a population (b), or annually kills all individuals in a subset of vulnerable patches (c). Colors and contours both denote population growth relative to ideal conditions without advection or disturbance. Life history parameters reflect our *P. popeii* study population (see Table 1).

### Case study: habitat distribution of P. popeii

To resolve *P. popeii* habitat distribution in the Rio Grande River, we surveyed 150 km of the river around Laredo (Texas; Fig. 4) in 2011-2012. Previously in this system Karatayev et al. (2018) found *P. popeii* almost exclusively in narrow gaps beneath large sandstone rocks resting atop bedrock. In this study we first searched for these *P. popeii* habitats, which were found in discrete sections (“patches”) along the river; by sampling from an airboat at low water allowed us to easily detect all patches. For each habitat patch, we estimated patch area, mussel density using 3-15 0.25 m^2^ randomly placed quadrats or detailed area searches covering 4-35 m^2^ (depending on bed area), and measured the length of all mussels found. We then tested for significant variation in *P. popeii* density, abundance, and mean mussel length across these habitat patches using an analysis of variance.

**Figure 4.**
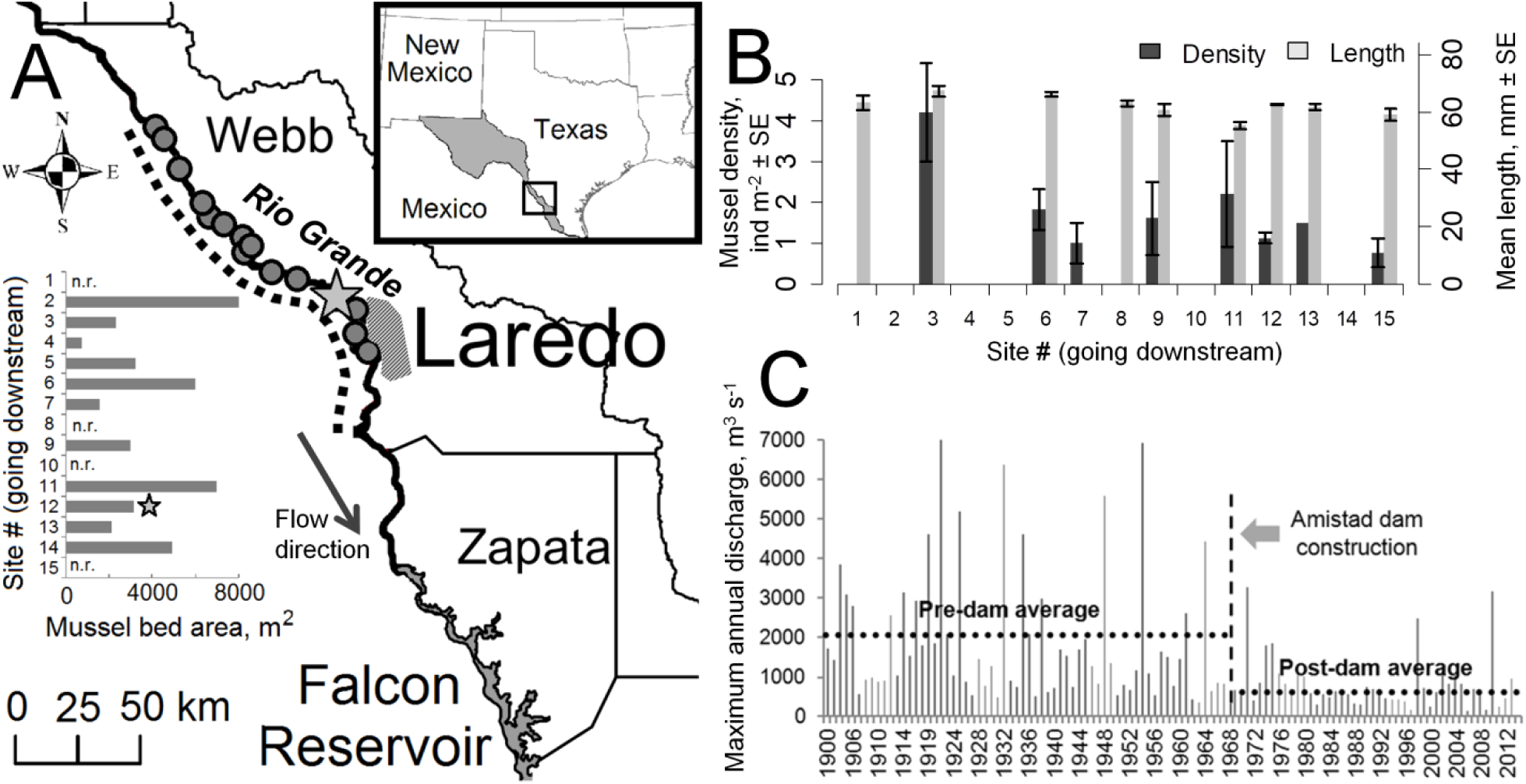
Distribution of *P. popeii* habitats (gray points on map) in 150 km section of the Rio Grande surveyed (a, dotted line), distribution of habitat patch areas (a, inset bar graph), variation in mussel density and mean size across habitats (b), and historic levels of peak annual discharge on the Rio Grande near Laredo, TX before and after the Amistad dam construction (c, maxima across daily readings; IBWC 2016). In (a), starred habitat denotes the mark-recapture site, the hashed area denotes the city limits of Laredo, and “n.r.” denotes “not recorded”. In (b) absence of bars for some sites indicates density or length measurements were not recorded.

### Case study: demography and spatial dynamics of P. popeii

To quantify *P. popeii* demography and spatial dynamics we conducted an intensive mark-recapture study at the La Bota Ranch site in Laredo in 2011-2014. Our Barker robust design approach annually sampled the population on three consecutive days in February-April to quantify and account for detection levels in our abundance and demographic estimates (White & Burnham 1999). This approach incorporates dead as well as live recaptures to estimate mortality separately from permanent downstream emigration. As live mussels could not burrow to avoid detection or secure themselves to counter strong currents, we assumed equal probabilities of detection and advective dispersal among live and dead individuals. We compared 32 parallel models that assumed constant, year-specific, and/or mussel length-specific survival and capture probabilities. To test for advective dispersal specifically, we compared models with versus without emigration. We obtained aggregate demographic estimates by averaging across estimates from all models, weighted by each model’s Akaike Information Criterion score. We then estimated immigration as the increases in abundance not explained by year-specific mortality, recruitment, emigration, or detection estimates. To test whether our analysis could produce plausible survival estimates without emigration, we also evaluated our best-fitting model (S4, Table 2) with emigration fixed at zero. For detailed mark-recapture methods see Appendix S4.

**Table 2.** Estimated population demography, abundance, and annual maximum discharge rates (across daily readings, IBWC 2016).

| Time Period | Encounter Probability | Frequency of recruits ($b_t/N_t$)<sup>a</sup> | Estimated abundance | Maximum Discharge ($m^3 s^{-1}$) | Emigration | Resident Survival | Frequency of Immigrants ($I_{t-1}/N_t$) |
| --- | --- | --- | --- | --- | --- | --- | --- |
| <u>March 8-11, 2011</u> | 0.11±0.02 | 0.03 | 986±145 |  |  |  |  |
| 2011-12 |  |  |  | 235 | 0.160±0.099 | 0.88±0.05 | 0.710 |
| <u>March 19-21, 2012</u> | 0.12±0.01 | 0.02 | 1463±134 |  |  |  |  |
| 2012-13 |  |  |  | 451 | 0.441±0.060 | 0.93±0.03 | 0.284 |
| <u>May 1-3, 2013</u> | 0.12±0.01 | 0.03 | 1178±122 |  |  |  |  |
| 2013-14 |  |  |  | 945 | 0.509±0.063 | 0.96±0.02 | 0.275 |
| <u>April 1-3, 2014</u> | 0.17±0.02 | 0.04 | 884±82 |  |  |  |  |

<sup>a</sup>Recruitment $b_t$ was estimated as the abundance of age-3 mussels in each year.
All uncertainties denote standard errors

To independently validate our model-fitted survival and emigration estimates, we additionally measured mean mussel longevity based on the size distribution of dead mussels and the magnitude of interannual variation in population size and age structure, which may reflect high levels of downstream migration. For this, we determined a size-age relationship (i.e., growth rate K and size at zero growth L_∞_) by fitting a von Bertalanffy growth model using least squares regression to growth observed in recaptured individuals and growth estimates based on early growth increments evident as imprints in mussel shells (Fig. 5b). Imprint-based growth estimates agreed well with observed growth, and were excluded in the few cases where imprints were obscure. In rare cases when length exceeded L_∞_ we assigned the minimum predicted age at L_∞_. We then estimated age distributions using the relation Age = – K^-1^log(1 – L_∞_^-1^Length), and used the mean age of dead *P. popeii* (i.e., longevity) estimate survival as exp(–Longevity^-1^). Finally, we used two-sample Cramer von Mises tests to determine the significance of interannual variations in size structure among all individuals caught (Fig. 5a) and individuals newly marked each year. Using this test statistic, we also compared the magnitude of year-to-year changes in *P. popeii* size structure in our study *vs.* a long-term study in the Black River, NM (Inoue et al. 2014). In all cases, we corrected size distributions for length-dependent detection estimated in our mark-recapture analysis (Fig. 5a, inset).

**Figure 5.**
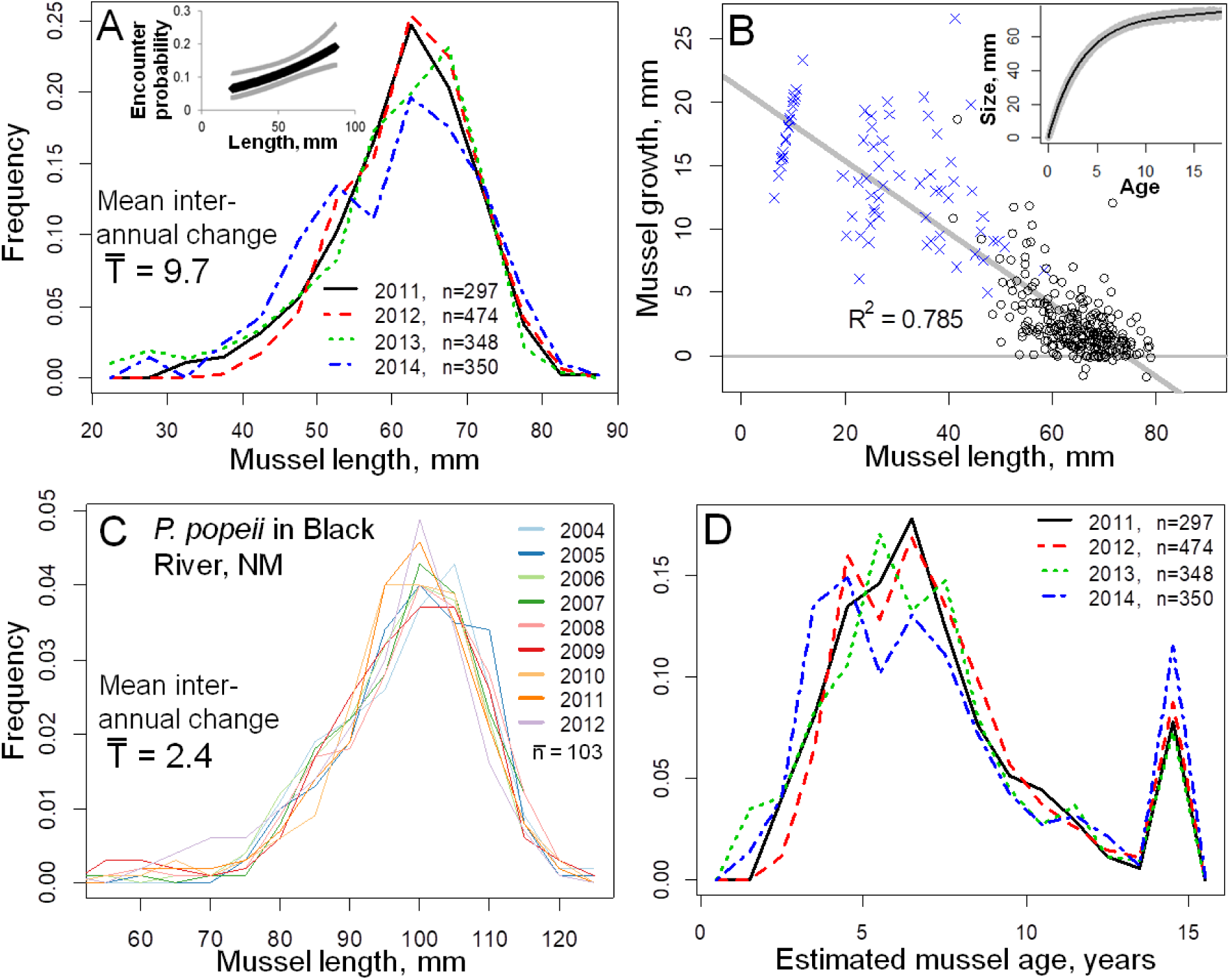
Large inter-annual variations of frequency distributions in size (a) and growth-based estimates of age (d) classes compared to variation in size-frequency distributions over a long-term study of P. *popeii* in the Black River, NM (c). Panel (b) denotes estimates of mussel age from length based on annual growth observed during study (circles) and juvenile growth estimated from shells (crosses), and the inset shows size-at-age (black) ± 1 standard error (gray region) based on uncertainties in K and L∞ (slope and x-intercept of regression in b, respectively). Numbers in legend of (a, c, d) indicate sample sizes used in size and age distributions (all mussels captured in a year). Inset in (a) shows the model averaged relation between mussel length and detection probabilities (gray lines indicate standard error) used to adjust the size-frequency distributions. 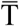 in (a) and (c) is the mean Cramer-von Mises test statistic for difference in size distributions computed over consecutive years. The peak at 15 years in (d) represents individuals longer than L∞, which were ≥15 years old and for which age could not be estimated.

## Results

### Washout effects on persistence across taxa

Washout effects reduced population growth for every life history examined (Fig. 2), including models that explicitly accounted for juvenile and adult stages (Appendix S1). In pristine habitats with all patches intact, advective dispersal alone reduced population growth only slightly via the added dispersal mortality and washout of adults from the patch furthest downstream. In degraded rivers with little advective dispersal, habitat loss alone weakly reduced persistence of taxa that cannot avoid poor habitats during biological dispersal, for instance winged insects or mussel larvae parasitizing fish. However, when intense flows and habitat loss co-occur, advection throughout the system transports individuals into degraded areas, producing the washout effect that reduced survival of affected stages and therefore overall population growth. Overall, stronger washout effects occurred in taxa vulnerable to advection throughout the majority of their lifespan (e.g., mussels and insects), particularly in longer-lived taxa. Washout effects also greatly affected fishes where only juvenile stages experience advective dispersal because young individuals are disproportionally more abundant in recovering populations, for instance short-lived fish with high per capita recruitment. The strength of washout effects was consistent across large (50%) deviations from estimated parameters, and depended predominantly on advective dispersal frequency, habitat loss, and per capita recruitment rather than poorly resolved dispersal distances (Fig. 2).

### Washout effects across habitat loss scenarios

Qualitatively different forms of habitat loss can lead to strong washout effects that reduce population persistence. In addition to long-term, small-scale habitat loss (Fig. 2; 3a), washout effects also occurred when long-term and large-scale habitat loss constrains populations to contiguous but increasingly small habitat areas (Fig. 3b). In this case, advection moves a higher fraction of the remnant population into inhospitable downstream areas as the number of habitat patches declines. Analogously, in metapopulations where temporary habitat loss is more likely in some river stretches, washout effects reduce persistence by moving individuals from better to less protected habitats, such as shallower pools vulnerable to drought (Fig. 3c).

### Habitat and population distribution of P. popeii

Along the surveyed 150 km river stretch we identified 14 suitable habitat patches. Total area of all patches corresponded to approximately 3% of the surveyed riverbed, and individual patches were separated by 1-5 km river stretches with a flat bedrock substrate where no *P. popeii* occurred. Habitat patches varied in area (3,589±785 m^2^, mean ± SE here and throughout; Fig. 4) but had very similar substrates, water depth, and hydrologic features (i.e., discharge flow rate and water velocity). All habitat patches contained live mussels at consistent densities (1.77±0.38 m^-2^, F=1.07, P=0.44, df=6, one way ANOVA) and similar mean length (P=0.21, F=1.62, df=7), suggesting that demographic rates at our study site were representative of the overall population. The furthest downstream inhabited patch was immediately upstream from the Laredo Sewage Plant wastewater discharge. In the 40 km surveyed area downstream of this site, rocky habitat patches with apparently suitable substrates contained no live *P. popeii*, likely due to organic pollution causing prominent biofilm layers.

### Estimates of advective dispersal in P. popeii

At local scales, several lines of evidence indicated high levels of adult mussel dispersal in our intensively studied mussel bed at La Bota. First, all mark-recapture models which received any AICc weight assumed high levels of emigration after accounting for interannual variation in survival and capture probabilities (Table 2, S4). Annual survival estimated by these models was consistent with our (conservative) survival estimates based on growth rates and the sizes of dead mussels (0.87±0.07, fraction surviving). By contrast, when assuming the death of all emigrants our best-fitting model estimated much lower survival (0.74±0.05, fraction surviving) and longevity (3.5±0.8 years). Second, high magnitude and variability in net migration explains the strong, significant interannual changes observed in mussel abundance (Table 2) given consistently low recruitment (i.e., frequency of individuals age ≤3), survival, and capture levels throughout our study. Finally, very strong interannual variation in *P. popeii* population age and size structure (Fig. 5a, d) indicated immigration from upstream habitats rather than a demographically closed, gradually aging local population. Size structure of mussels newly marked each year differed significantly among sampling events spanning two years with the highest maximum discharge levels (2011-12: T=2.9, p=0.63; 2012-13: T=22.6, p=0.008; 2013-14: T=16.0, p=0.04, Cramer von Mises tests, sample sizes 297-474). For the whole population, inter-annual variation in size structure was 3-fold higher compared to a 9-year study of *P. popeii* in the Black River, where peak discharges are three orders of magnitude lower (Fig. 5c; Inoue et al. 2014).

## Discussion

In heavily impacted systems where extensive habitat loss coincides with transformed flow regimes, restoring historic peak river flows alone may have detrimental impacts by intensifying washout of threatened populations out of the remaining suitable habitats. This unintended consequence can greatly accelerate local extirpation across taxa with distinct life histories (Fig. 1, 2). Washout can occur regardless of whether habitat loss manifests at localized scales throughout the range of a population (Fig. 3a), at large scales by isolating subpopulations to small river stretches (Fig. 3b), or temporarily as short-term disturbances (Fig. 3c).

Avoiding the adverse effects of restoring natural peak flows requires quantifying advective dispersal and, where it greatly affects populations, restoring habitats in conjunction with the flow regime. Our empirical study is the first to quantify advective dispersal in adult stages of freshwater mussels, a particularly threatened taxon. High levels of advection found here (16-51% of population annually, Table 2) reveal that mussel populations might have strong spatial dynamics and a high vulnerability to washout effects (Fig. 2). Across taxa, restoration efforts could alleviate washout by creating or maintaining long, contiguous stretches of high-quality habitats wherein washout effects occur only in downstream patches (Fig. 3b). The required habitat area increases with the magnitude of advective dispersal (Fig. 3b), and patch size will likely depend on the imperiled species (see *Methods* and below). Vice versa, conservation efforts may fail to prevent local extinction when protecting even abundant subpopulations isolated in small remnant habitats as elevated peak flows increase advection.

### Evidence for and detection of advective dispersal under peak flows

A common theme of emerging empirical studies across taxa is that advective dispersal of vulnerable life stages can be common and occurs primarily under peak river flows. Most studies explicitly tracking flow-driven movement resolve only localized “behavioral” drift (e.g., <200m, Brittain & Eikeland 1988) at low discharge levels. However, field studies spanning flood events find mass advection of aquatic insects (Brittain & Eikeland 1988; Gibbins et al. 2007), juvenile fish (Lechner et al. 2016), and molluscs (Kappes & Haase, 2012). In turn, peak flow conditions can also determine the spatial distribution of organisms (Brittain & Eikeland 1988): for instance, Gangloff and Feminella (2007) found that bankfull shear stress predominantly governed spatial mussel distribution across 8 streams. Finally, site-specific studies find lower retention of marked individuals in years of higher discharge (Villela et al. 2004), a pattern also seen in our results (Table 2) and restoration efforts in which floods washed away reintroduced mussels out of entire habitat patches (Ahlstedt 1980; Layzer & Gordon 1993). These results suggest a strong influence of peak flows on the spatial dynamics of many organisms, paralleling foundational ideas in hydrology that spatial dynamics of sediments and organic matter predominantly depend on peak discharge (Yarnell et al. 2015).

Field studies also find direct evidence for advective dispersal over long distances in relatively sessile organisms moved predominantly by river currents. Spatial surveys often find individuals established in previously unoccupied habitats following floods (gastropods: Rosa et al. 2014; mussels: Hastie et al. 2001; Sousa et al. 2012). Surveys in systems with concurrent mark-recapture studies have also found adult mussels living long distances (1-10km) downstream from tagging sites (Balfour & Smock, 1995; Welte, Dunn, Alderman, personal communication). In single-site demographic studies that do not consider migration, the presence of advective dispersal might also explain anomalously low mussel survival estimates (0.50, Villela et al. 2004; 0.49±0.04, Matter et al. 2013; 0.58±0.06, our estimates ignoring emigration) that contradict the persistent, adult-dominated populations observed in these systems. In many such rivers where few habitats offer permanent refuge from peak flows, our results suggest that (1) advection can move a large fraction of adult mussels among habitat patches, and (2) this spatial dynamic does not greatly reduce population persistence in pristine systems (e.g., 4%, Fig. 3a). In the Rio Grande, frequent advective dispersal among separate habitat patches appears likely given that we found *P. popeii* exclusively under large rocks, substrates offering the strongest resistance to current in the bedrock-lined riverbed. This reflects previous field studies and experiments demonstrating that peak flows can concentrate mussels in areas with weaker river currents (Gangloff and Feminella 2007), move adult mussels out of these habitats (Strayer 1999), and transport them to downstream flow refugia (Daraio et al. 2012).

Demography-based approaches could estimate advective dispersal beyond specific sessile taxa, but may require intensive studies at limited spatial and temporal scales. Advective dispersal may be directly measured by tracking tagged or isotopically marked individuals (Hershey et al. 1993; Bilton et al. 2001). Such efforts can be hampered by low detection rates (e.g., mussels, Villela et al. 2004; Table 2) or high mortality of organisms affected by advection (e.g., juvenile fish, Franzin & Harbicht 1992). An alternative approach is to quantify mortality separately from dispersal by accounting for recovery rates of dead individuals in mark-recapture studies (as we have done here) or estimating mortality from changes in population age or size structure. Advective dispersal is then the difference between observed changes in abundance and those predicted by mortality and recruitment alone. Unfortunately, detailed demographic studies span limited spatiotemporal scales and might underestimate dispersal when they fail to span high-flow areas or infrequent (e.g., decadal) peak flows that have disproportionally strong population effects, especially on long-lived species (Strayer 2008).

We suggest that resolving the magnitude of advective dispersal and resulting washout effects may be best achieved using approaches which synthesize hydrodynamic studies and field surveys. Hydrodynamic models and experiments are increasingly used to resolve the potential for flow-driven movement of organisms across taxa and observed flow rates in a given system (Strayer 1999; Gibbins et al. 2007). When this potential for advective dispersal is strong, hydrodynamic models can predict local peak flow intensity across occupied and unoccupied habitats using broad substrate and streambed characteristics (Morales et al. 2006; Daraio et al. 2012). Evaluating the extent to which peak flow characteristics *vs.* other habitat requirements explain the spatial distribution of vulnerable populations (e.g., Gangloff & Feminella 2007; Maloney et al. 2012) could then estimate realized advective dispersal. A particular strength of this approach is the potential to predict advection across long river stretches, historic floods, and the target flow regime. Field surveys comparing population distribution before *vs.* immediately after peak flow events could further supplement efforts to quantify advective dispersal. When they identify intense advective dispersal, joint modeling and empirical approaches can directly prioritize habitat restoration in river stretches with the greatest potential for washout effects.

### When restoring peak flows might reduce local persistence

Restoring peak river flows might reduce local persistence when (1) advection affects a large portion of a population and (2) extensive habitat degradation spans sufficiently large spatial scales. The prevalence of advective dispersal in a population depends on the relative abundance of life stages vulnerable to drift under peak flows and the frequency of peak flow events (Fig. 2). However, in long-lived species even infrequent peak flow events on generational timescales can lead to washout. Although strongest in sessile taxa like mussels, washout effects might reduce persistence even when adults can actively move long distances because the structure of recovering populations is generally skewed towards younger, more vulnerable stages.

Washout effects in a given taxa require habitat loss that spans spatial scales (e.g., river length) greater than the distance individuals can freely move, in which case organisms displaced by flow into degraded areas cannot reach suitable habitats before dying. We implicitly assume this by modeling habitat patches among which fish larvae, insects, and molluscs disperse only annually and patch-scale habitat loss. This approach allowed us to compare washout effects among life histories independently from context-specific drivers of habitat loss. However, within a given system organisms affected by advection might greatly differ in movement capacity and therefore in their sensitivity to habitat loss. For instance, even fine-scale (∼100m) habitat loss can produce washout effects in molluscs that actively move only short distances. In contrast, fish and insect larvae which electively re-enter river currents and drift to more suitable downstream habitats (Townsend 1989) may only be affected by washout into larger (e.g., 1-5 km) sections of degraded habitats. These areas might include river stretches with adverse food, predation, or abiotic regimes such as reservoirs (Hofer & Kirchofer 1996), river channels with low substrate complexity, or areas of intense pollution.

When restoring peak flows in heavily degraded systems, two exceptions might reduce the capacity for washout effects. First, restoring habitats by reducing hypoxia and substrate siltation is a common goal of restoring river hydrology, and restoring peak flows in particular (Yarnell et al. 2015). If flow restoration alone can improve habitat conditions quickly relative to the annual rate of advective dispersal, it may have a positive net effect on population persistence. Second, extensive habitat loss might have limited impacts on persistence if sufficiently large, contiguous habitats remain. For instance, despite the loss of *P. popeii* from 75% of its historical range in the Rio Grande, relatively large sections of suitable habitats remain (∼15-28 patches, Karatayev et al. 2018). However, such constricted populations are particularly vulnerable to further temporary or long-term habitat loss and fragmentation, for example due to dessication during droughts or organic pollution that occur in our study area (Karatayev et al. 2018). In these cases, flow restoration places greater importance on protecting existing habitats. Finally, we note that two potential benefits of increased advective dispersal, habitat recolonization and genetic connectivity, might be redundant with naturally evolved forms of dispersal. In all of our temporary habitat loss simulations washout reversed the benefits of recolonization (Fig. 3c, C.1).

### Conclusions

Natural resource managers increasingly rely on environmental flow regulation to restore ecosystem processes and native populations. We highlight a potential to accelerate rather than reverse species loss when restoring peak flows to natural levels in a system that has also lost a large proportion of suitable habitat for a particular species of concern. This adverse impact is due to advective transport of the species from remnant suitable habitats into degraded areas. We suggest integrating hydrodynamic studies and field surveys to detect both the potential for advective dispersal in specific systems and its population consequences. Where this potential is strong, we recommend a multi-faceted conservation approach concomitantly restoring flow regimes and long, contiguous stretches of natural habitats.

## Supporting Information

Effects of stage structure (Appendix S1), parameterization of spatial population models (Appendix S2), washout effects under temporary habitat loss (Appendix S3), and detailed mark-recapture methods (Appendix S4) are available online. The authors are solely responsible for the content and functionality of these materials. Queries (other than absence of the material) should be directed to the corresponding author.

## Appendices

### Appendix A: Effects of stage structure

By adding a delay between recruitment of new individuals and when they reproduce, during which immature “juvenile” stages are susceptible to mortality, may affect our results regarding the effects of advective dispersal. Here we (1) reproduce repeat our main analysis of the interactive effect of advective dispersal and habitat loss on population growth λ and (2) compare the sensitivity of λ to model parameters in the structured *vs.* unstructured model versions.

#### Model setup

For simplicity, we structure the population with two stages, rather than continuous ages, though the qualitative effects are the same for models with three or more stages (results not shown).

**Figure A.1.**
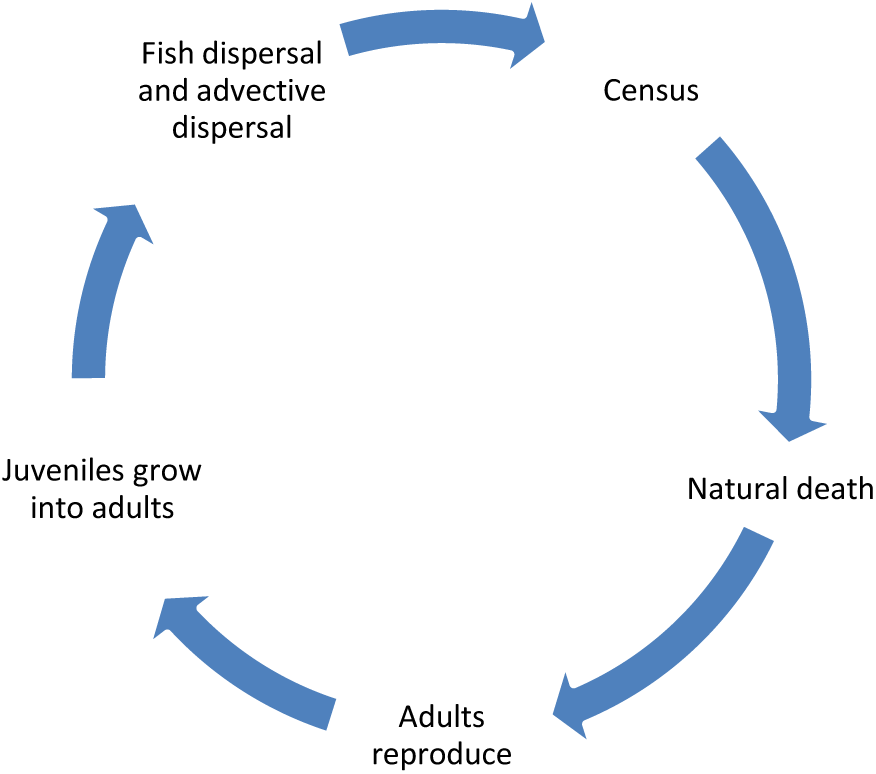
Life cycle of the stage-structured mussel population

In our model juveniles who cannot reproduce and sexually mature adults are distributed across L patches. The state of the population is given by the matrix

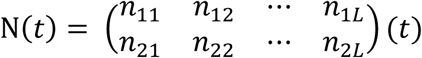

where *n*_*ij*_(*t*) is the abundance of mussels in patch i, stage j in year t. This matrix can be rewritten as a population vector arranged by patches, allowing us to explore population dynamics using projection matrices.

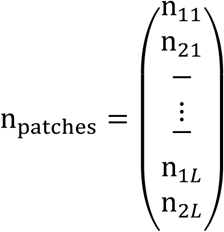

We calculated the mussel abundance next year (N(*t*+1)) using a vec-permutation matrix approach (Caswell 2012). Under our assumption that demographic changes happen before dispersal (Fig. A.1), the projection matrix including both demographic and dispersal processes can be written as a combination of processes affecting the survival and advective dispersal of individuals already in the population (Ũ) and the abundance and spatial distribution of newly added recruits 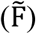:

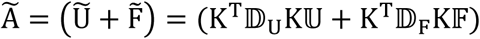

Mussels in patch i who don’t die of natural mortality (1 − *d*) first go through reproduction (*R* for adults) and stage transition (*g* for juveniles) Thus, the stage projection matrix A_*i*_ can be decomposed into

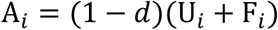

where

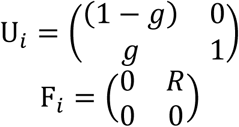

All newly produced offspring are subsequently dispersed by fish to *n* neighboring patches upstream of patch i and *n* neighboring patches downstream of patch i, and pre-existing individuals (juveniles and adults) are removed by flow (proportion *ρ*) to *m* neighboring patches downstream of patch i, some of which may die during dispersal (proportion *µ*). For example, dispersal can be described by the following two matrices for L = 5 patches where m = 1, and n = 2:

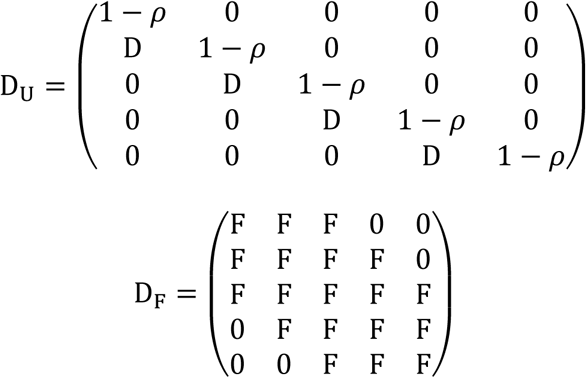

Where .D = *ρ*(1– *μ*)/*m*, and F=1/(2*n*+1).

In the final expression of projection matrix Ã, matrices 𝕌, 𝔽,𝔻_U_ and 𝔻_F_ are block-diagonal matrices constructed by calculating the Kronecker product of matrices U_*i*_,F_*i*_,D_U_,D_F_, and corresponding identity matrices, respectively. The vec-permutation matrices K and K^T^ are used to convert the population vector by patches into population vector arranged by stages or vice versa, respectively.

We used the same natural mortality for different stages (mark-recapture data on >20mm -age 2 -individuals do not suggest a strong effect of size on natural, authors’ unpublished data). Lacking data on age at maturity for *P. popeii*, we chose a moderate estimate of g = 0.6 corresponding to a mean age at maturity of 2. All other parameters were retained from the original model (Table 1). In all cases, we repeated of main analysis by using two versions of the model: one unstructured with 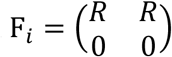 and *g*=0, and one incorporating stage structure with 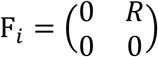 and *g*=0.6.

#### Effects of stage structure on the interaction between advective dispersal and habitat loss

In the stage structured population model, overall population growth is always lower than in the unstructured model (Fig. A.2, A.3) because natural and dispersal-associated mortalities among juveniles effectively reduce adult fecundity R. Increasing the frequency of invalid patches (where adult survival is zero) (Fig. A.2, left panel) and decreasing number of adjacent patches (Fig. A.2, right panel) have the same qualitative effects on stage-structured and unstructured model. However, the difference between stage-structured and unstructured model shrinks when most patches are lost or the small domain contains very few patches (less than 5). In such cases of very small habitat domains, loss of recruits due to settlement into invalid patches is much greater than the effective reductions in adult fecundity by juvenile mortality.

Similarly, when temporary disturbances are happening to randomly selected patches, the effects of advective dispersal on the growth of stage-structured populations is qualitatively the same as in the unstructured case (Fig. A.3). Again, since the mortality of immature individuals again effectively reduces adult fecundity, growth of structured populations are consistently lower.

**Figure A.2:**
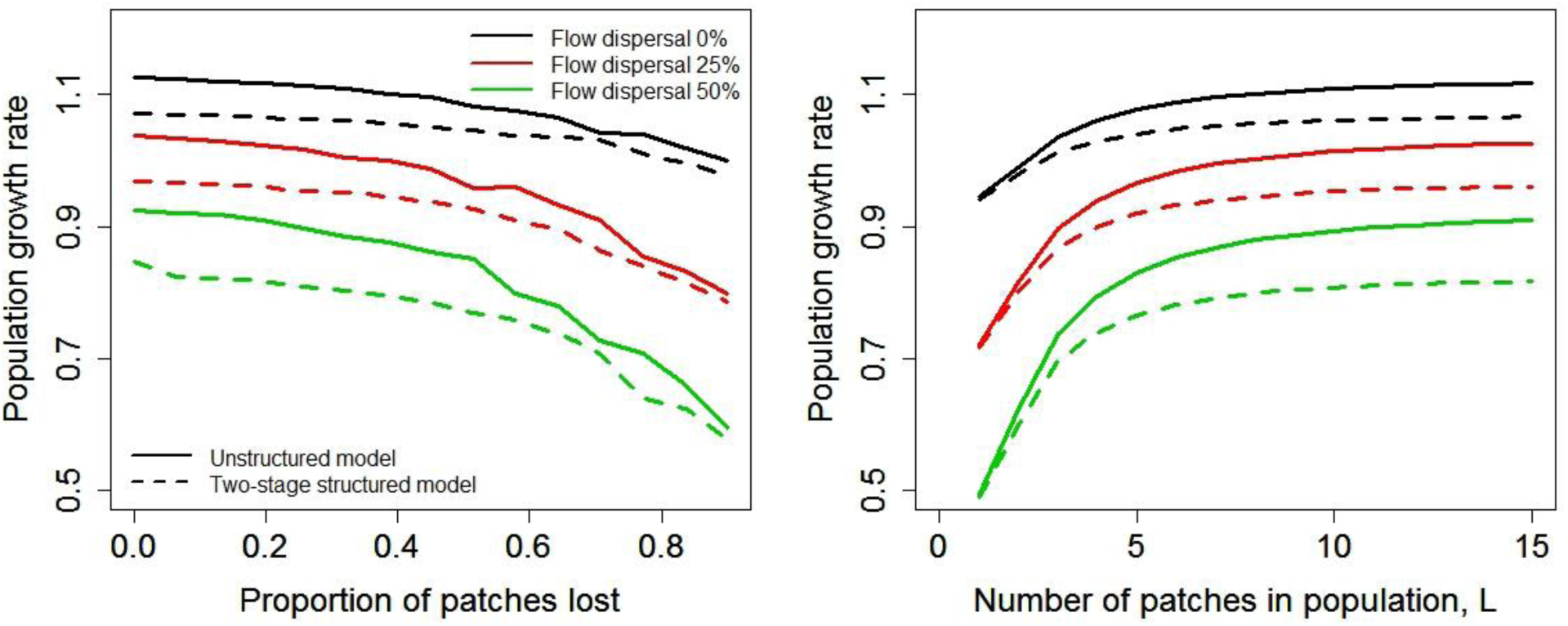
Advective dispersal amplifies the decrease in population growth *λ* with increasing levels of permanent habitat loss (left panel), and as the number of patches remaining in a population decreases (right panel). Stage structure has a small negative effect on population growth *λ*, but also reduces the decreases in population growth *λ* caused by increasing proportion of permanent patch loss or decreasing number of patches in a small domain.

**Figure A.3.**
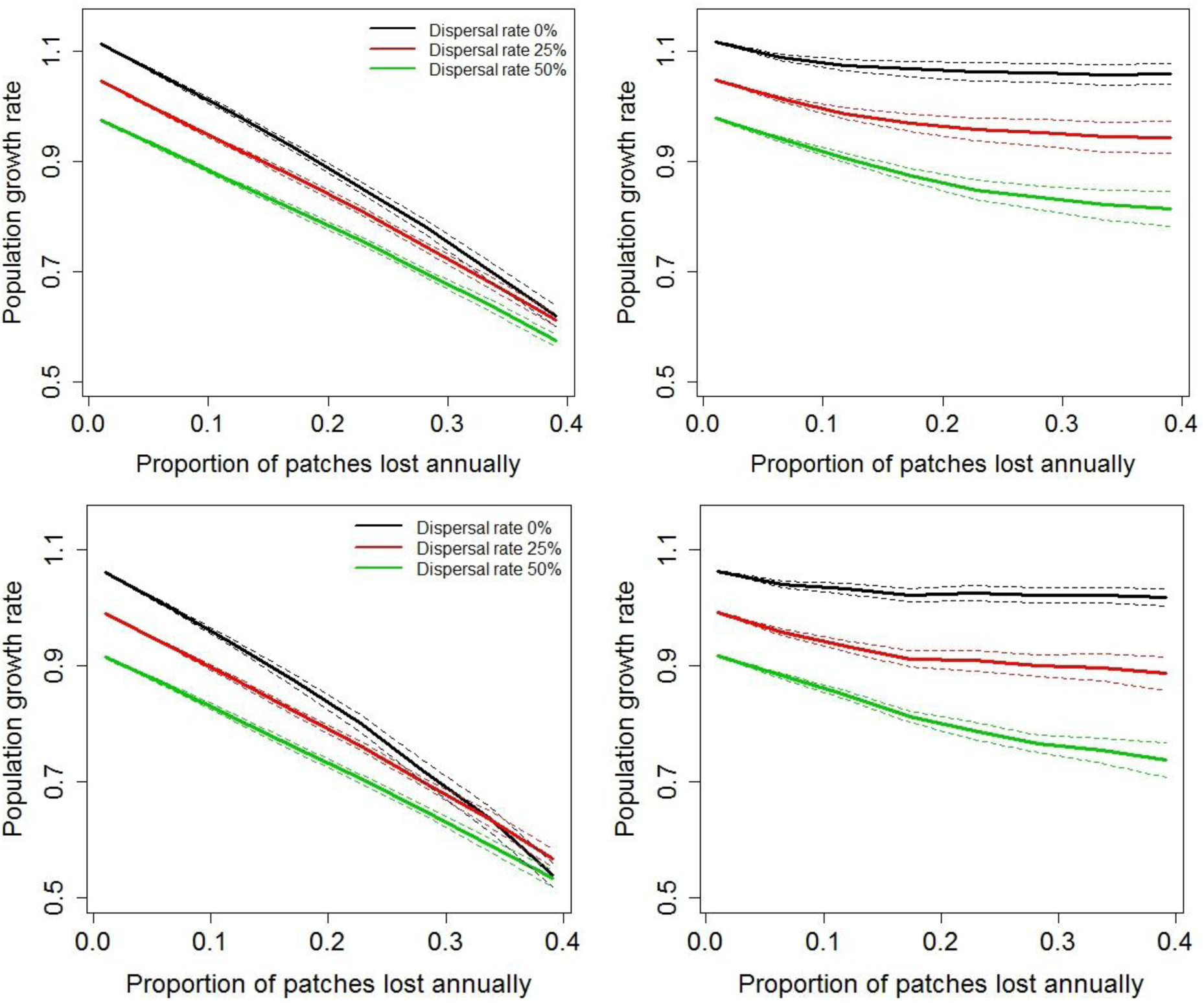
Analysis in main text reproduced without (top panels) and with (bottom panels) stage structure in the population. In both cases, advective dispersal (colored lines) has no effect on the decrease in population growth λ with increasing disturbance frequency when all patches are disturbed with equal frequency (A), but promotes the decline in growth under increasing disturbance when some habitat patches are disturbed less often (B). Here, disturbance frequency is the proportion of 100 patches from which the population is extirpated annually (i.e., all adults are killed by drought) (A) or the proportion of 70 vulnerable patches annually becoming extirpated, with the remaining 30 invulnerable to disturbance. Dotted lines denote variability in λ values due to different spatial distribution of disturbances along the river length.

## Appendix B. Parameterization of spatial population models

In comparing washout effects across life histories, we use a level of 85% habitat loss that is broadly consistent across studies of range contraction in fish (Cahaba shiner, 83%; Rio Grande fishes, 93%, Malloy et al. 2013; *Oncorhynchus mykiss* in California, 40%, Boughton et al. 2005), freshwater mussels (*P. popeii* in the Rio Grande, 75%, Karatayev et al. 2012), and other invertebrates such as shrimp (*Macrobrachium* sp., 66%, Bowles et al. 2000). Given that range contraction might arise only partly due to habitat loss, however, context-specific models may be required to determine whether extensive range contraction is associated with a large potential for washout effects.

Throughout, we point out that poorly resolved estimates of maturation/emergence, reproduction, and dispersal had very weak impacts on the effects of washout in our sensitivity analysis (except for reproduction r for fishes Fig. 1).

### Freshwater mussels

For our simulations, we used the average mortality (d) and probability of emigration (ρ) measured in our 4-year study. Given that patches in our studied population are relatively short (100-500m) but situated at 1-5km intervals, we set fish dispersal distance n=2 patches=∼14km. Although this is a generous estimate for ecological hosts of *P. popeii* in the Rio Grande (Caldwell pers. comm.), fish hosts of other species can move mussel larvae across much larger distances (Woolnough et al. 2009). Lacking data on distances of advective mussel dispersal, we chose a reasonable estimate of m=1. Although true values for flow and fish-driven mussel dispersal will require much effort to estimate (and dispersal distances across host fish species are particularly variable; Woolnough et al. 2009), our results are qualitatively the same for different levels of n, m >0. Also, whereas dispersal may not be evenly distributed across space and biased towards upstream or downstream sites, these simplifications have no effect on our results due to our assumption that habitat loss occurs randomly across habitat patches in the river. The number of offspring surviving to sexual maturity generated annually by each adult (r) is difficult to measure in many systems; we estimated a mean mussel age in our population of ∼8 years (based on our growth and size structure data), and assuming a maturation age of 2 years and that 50% of individuals are females, we chose r=0.4 (i.e., on average females throughout their lifetime produce 4.8 offspring that survive to maturity). Finally, we chose a moderate level of adult mortality during advective dispersal μ=0.3, although our qualitative results are robust to other values of this parameter.

In parameterizing the model for a long-lived mussel species, *Margaritifera margaritifera*, we assume a lower recruitment level (r=0.125, which yields λ=1.1) to account for the older age at which individuals mature. We derive mortality from the maximum age A_max_ reported for this species by Hastie (2000). As a precise definition of A_max_ is not given, we assume it corresponds to an age at which 1% of individuals in a given cohort remain; thus, we assume a constant mortality (d) and calculate it as: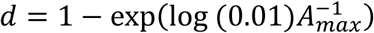. Finally, to facilitate comparison with our results for *P. popeii* and given the paucity of information on these parameters, we retain the same values of m, n, ρ, and µ; again, we examine the sensitivity of these parameter choices below.

### Aquatic insects

This model tracks the spatiotemporal dynamics of larvae in a recovering population of hydropsychid caddisflies. Although the amount of time required by larvae to reach maturity and emerge varies across populations and species, hydropsychids are typically univoltine, and so we assume that most individuals emerge within a single year (γ=0.8). Prior to emergence (i.e., through the larval and pupal stages), based on a range of larval and pupal survival estimates (Merritt and Cummins 1996), we assume a moderate level of mortality (d=0.5) which corresponds to an average longevity of 1.5 years for larvae which do not emerge. Note that the comparatively low survival of adult (winged) insects (Merritt and Cummins 1996) is implicitly accounted for in r. After emergence, dispersal in hydropsychids has been estimated at 0.5-1.5km (Kovats et al. 1996). In this model, we assume habitat patches are 0.5-1km long, so approximately one-third of dispersing adults reach an adjacent downstream or upstream patch (n=0.35). For advective dispersal, we assume that a proportion ρ=0.5 of both newly recruited individuals and those remaining from previous years are annually subjected to advective dispersal and are moved downstream by up to m=2 patches.

### Fish

In this model we assume distinct patches of suitable habitats are 1-5km long and parameterize the model for two European fishes commonly found to dominate the assemblages of drifting fish larvae, *Alburnus alburnus*, and the longer-lived *Abramis brama* (Zitek et al. 2004). Using maximum age estimates in Blanck et al. (2007), we estimate mortality d following the relation used above for long-lived mussels. Because larvae spend weeks to months in developmental stages which are vulnerable to or actively participate in advective dispersal, we conservatively set ρ=0.5 in this analysis. Finally, we assume that adults on average move n=5 habitat patches (1-25km in our model) away from the patch where they reproduced in the prior year, and that advective dispersal can also transport larvae over distances of up to m=4 patches, i.e. on average 4-8km downstream of the natal patch.

## Appendix C. Washout effects under temporary habitat loss with all patches vulnerable

When random, localized extirpations happen from temporary habitat loss, washout effects reduce population growth only slightly because in the long-term, habitat quality is identical in all patches (Fig. C.1a). However, when some habitats are always less vulnerable to local extirpation, washout effects arise again as advective dispersal continually transports individuals from better protected, undisturbed areas to patches where extirpations are more frequent (Fig. 3c). Thus, as with long-term habitat loss, advective dispersal in this case interacts with the frequency of local disturbance to reduce population persistence.

**Figure C.1.**
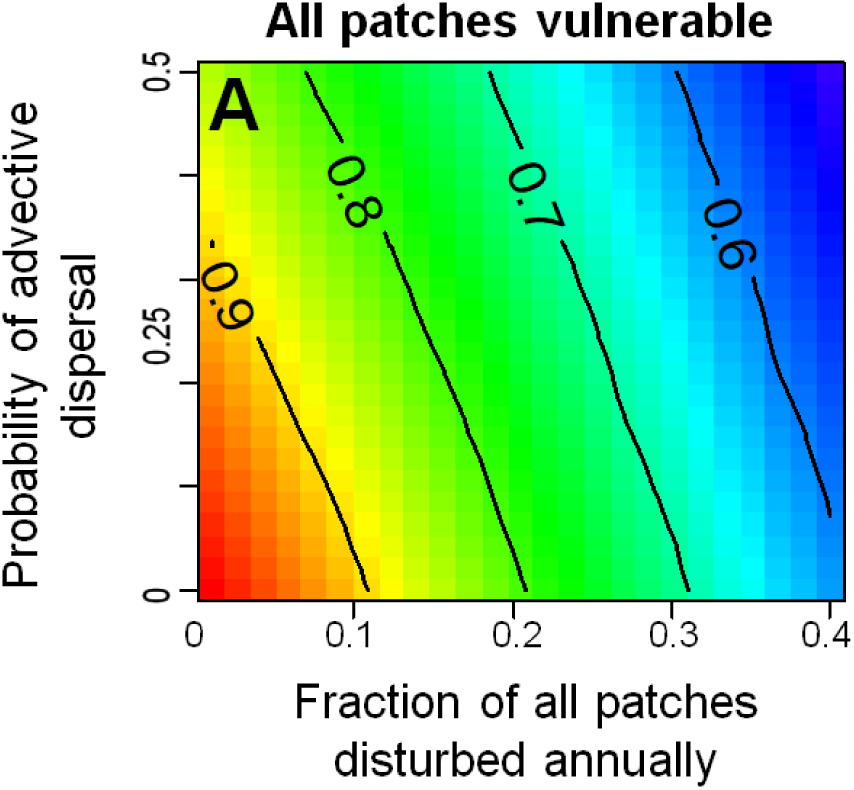
Effects of habitat loss and advective dispersal on population growth when disturbance-induced local extirpation is equally likely in all patches. Colors and contours denote population growth relative to ideal conditions without advection or disturbance. Parameters are fitted to our *P. popeii* study population (see Table 1).

## Appendix D. Detailed mark-recapture methods

Access to the site was gained from the US Border Patrol boat ramp and owner permission. Each year we located and marked the ca. 1,000 m^2^ mussel bed using GPS coordinates and surveyed the area on three consecutive days (21 person hours mean daily sampling effort), during which period we assumed the population was closed (i.e., no recruitment, mortality, or dispersal). We measured the length of all mussels and marked individuals captured for the first time throughout our study with numbered Floy laminated flex tags (1231 individuals total).

We estimated demographic parameters using a Barker robust design mark-recapture model and a Huggins Closed Captures sample design with live and dead recaptures using program MARK 8.2 (White and Burnham, 1999; Barker 1997). We adjusted our analysis to account for the minor differences in durations between interannual sampling events. We then constructed a parallel set of 32 models which compared parameter estimates (annual probabilities of survival S, capture p, and permanent emigration F) across models which assumed the parameters were constant or variable across years, and combinations of these. We also included models assuming an effect of size on survival and encounter probabilities. We tested for handling effects on survival by modifying our best-fitting model to account for difference among survival of newly vs. previously marked individuals. Finally, we quantified the number of immigrants *I*_*t*_ arriving between year *t* and *t*+1 using time-dependent model averaged estimates of abundance (*N*_*t*_), survival (*S*_*t*_), and emigration (*F*_*t*_) and estimates of recruit abundance (*b*_*t+1*_; number of age-3 mussels) based on the age distribution of live mussels using the relation

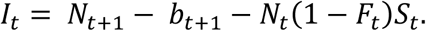

All mark-recapture models which received any AICc weight assumed high levels of emigration and interannual variability in survival and capture probabilities (Table D.1). Models with and without handling effects had equivalent performance (AICc score difference < 2, Burnham and Anderson, 2002). Derived estimates of immigration differed markedly from the estimated emigration in some years, as reflected in the large changes in abundance (Table 2).

**Table D.1.** The five best-fitting models from 34 candidate models in our Barker robust design analysis. S, F, and p denote survival, emigration, and recapture levels, t and L denote the dependence of these factors on year and size when marked, and h denotes an effect of handling on survival. Final row gives fit of our best-fitting model without emigration.

| Model | AICc | Delta AICc | AICc Weights | Model Likelihood | Number of Parameters |
| --- | --- | --- | --- | --- | --- |
| $S(t+L) F(t) p(t+L)$ | 6348.1 | 0 | 0.64 | 1 | 20 |
| $S(t+h+L) F(t) p(t+L)$ | 6350.3 | 2.25 | 0.21 | 0.32 | 22 |
| $S(t) F(t) p(t+L)$ | 6351.5 | 3.40 | 0.12 | 0.18 | 19 |
| $S(t+L) F(t) p(t)$ | 6355.5 | 7.46 | 0.02 | 0.02 | 19 |
| $S(t+L) F(t) p(t)$ | 6356.1 | 8.06 | 0.01 | 0.02 | 18 |
| $S(t+L) F=0 p(t+L)$ | 6416.5 | 68.5 | 0.00 | 0.00 | 17 |

